# Transcriptional landscapes and signaling pathways of chloroquine-treated Esophageal squamous cell carcinoma

**DOI:** 10.1101/2022.08.19.504517

**Authors:** Wei (David) Wang, Zhiwen Qian

## Abstract

Esophageal squamous cell carcinoma (ESCC) is one of the human malignancies worldwide, but the mechanism of ESCC development is still unclear. Chloroquine has the anti-tumor function by the inhibition of autophagy and thereby contributing to apoptosis. In our study, we analyzed the RNA-seq data of Chloroquine-treated ESCC cells and identified the transcriptional landscapes. We then used the gene enrichment methods such as KEGG and GO to further analyze the potential signaling pathways. In addition, we constructed the PPI network and Reactome map to further identified the biological processes. We identified the top two signaling pathways that were involved in the chloroquine-treated ESCC: Cell cycle and Glycerophospholipid metabolism. We identified the top ten interactive genes including ATM, CCNB1, FN1, CCT6A, VEGFA, PA2G4, CCT2, CDKN1A, BRIX1, and CDC20. Our study may provide new insights into the mechanisms for the Chloroquine-treated ESCC cells.

## Introduction

Esophageal cancer includes squamous cell carcinoma and adenocarcinoma^1^. Developing countries showed higher incidence rates, such as Southeast Asia, Eastern and Southern Africa along the Indian Ocean coast, and South America^2^. Among the regions, Asia is the region with the highest incidence of esophageal cancer^3^. However, China has the highest incidence in Asia, which accounts for about half of the world’s esophageal cancer incidence^4^. Nearly all the histology of esophageal cancer noted is squamous cell carcinoma^5^.

Chloroquine was used to treat malaria and was also used to treat autoimmune diseases^6^. In addition, chloroquine shows the effects through the weak-base lysosome-tropic feature^7^. When chloroquine goes into the lysosome, it becomes protonated due to the low pH, and accumulation of the protonated form of chloroquine within the lysosome leads to less acidic conditions and further results in decreased lysosomal function^8^. Recent findings showed that chloroquine is promising for cancer treatment. Several clinical trials have shown favorable effects of chloroquine as an effective anti-cancer drug. However, the precise mechanism is not clear^9^.

In this study, we analyzed the gene profiles of Chloroquine-treated ESCC cells by using the RNA sequence data. We further identified the key molecules and signaling pathways by performing the KEGG, GO, Reactome map, and PPI methods. Therefore, our results showed the potential mechanisms of Chloroquine-treated ESCC.

## Materials and Methods

### Microarray data

GEO database (http://www.ncbi.nlm.nih.gov/geo/) was used for downloading the following gene expression profile dataset: GSE209582. Illumina NextSeq 500 (Homo sapiens) (Temple University, Fels Institute, Whelan Lab, 3307 N Broad St. PAHB Rm 203, Philadelphia, PA19140, USA). The dataset was chosen based on the criteria as follows: Studies comparing control untreated ESCC cells with chloroquine (10µM) treated ESCC cells; Studies included gene expression profiles; Datasets were excluded if the study did not contain control groups; Datasets from other organisms were excluded.

### Differentially expressed genes (DEGs)

The gene expression matrix data in the GSE209582 dataset was normalized and processed through the R program as described^10-14^. Then, the empirical Bayesian algorithm in the “limma” package was applied and compared between patients with diseases and the healthy control population as controls with the detection thresholds of adjusted P-values <0.05 and |log2FC| >1.

### Gene intersection between DEGs

We intersected the DEGs screened by the “limma” package with the modular genes. The intersecting genes were analyzed for Kyoto Encyclopedia of Genes and Genomes (KEGG) and Gene Ontology (GO) enrichment using the “clusterprofiler” package in the R program. The results were then visualized using the “ggplot2” package.

### Protein-protein interaction (PPI) networks

The interacting genes were constructed using Protein-Protein Interaction Networks Functional Enrichment Analysis (STRING; http://string.db.org). The Molecular Complex Detection (MCODE) was used to analyze the PPI networks. This method predicts the PPI network and provides an insight into the biological mechanisms. The biological processes analyses were further performed by using Reactome (https://reactome.org/), and P<0.05 was considered significant.

## Results

### DEGs in chloroquine-treated ESCC

To make sure the functions of **chloroquine** on **ESCC**, we analyzed the RNA-seq data from the GEO database (GSE209582). We identified the 770 significantly changed genes (P < 0.01). The top increased and decreased genes were also indicated by the heatmap (Figure 1). We further presented the top ten significant genes in Table 1.

**TABLE 1.**

| **ENTREZ**<br>**GENE** | **Gene symbol** | **Fold-change** | **Regulation** |
| --- | --- | --- | --- |
| **TOP 10 DOWN-REGULATED DEGS** |  |  |  |
| **3008** | H1-4 | -3.427517651 | Down |
| **3304** | HSPA1B | -1.914306042 | Down |
| **3303** | HSPA1A | -1.899342169 | Down |
| **257313** | UTS2B | -1.864020827 | Down |
| **6415** | SELENOW | -1.781354979 | Down |
| **102723780** | FRG1FP | -1.768259817 | Down |
| **100288661** | SUB1P4 | -1.729852077 | Down |
| **100288077** | WTAPP1 | -1.618740966 | Down |
| **280636** | SELENOH | -1.611152382 | Down |
| **100418905** | NBEAP3 | -1.574041963 | Down |
| **TOP 10 UP-REGULATED DEGS** |  |  |  |
| **390874** | ONECUT3 | 4.169739817 | up |
| **50509** | COL5A3 | 3.247308337 | up |
| **1303** | COL12A1 | 3.217243699 | up |
| **26353** | HSPB8 | 2.951933773 | up |
| **337882** | KRTAP19-1 | 2.815013049 | up |
| **1002** | CDH4 | 2.448801995 | up |
| **114815** | SORCS1 | 2.424578489 | up |
| **84966** | IGSF21 | 2.415838872 | up |
| **8399** | PLA2G10 | 2.322082632 | up |
| **80726** | IQCN | 2.251183802 | up |

**Figure 1.**
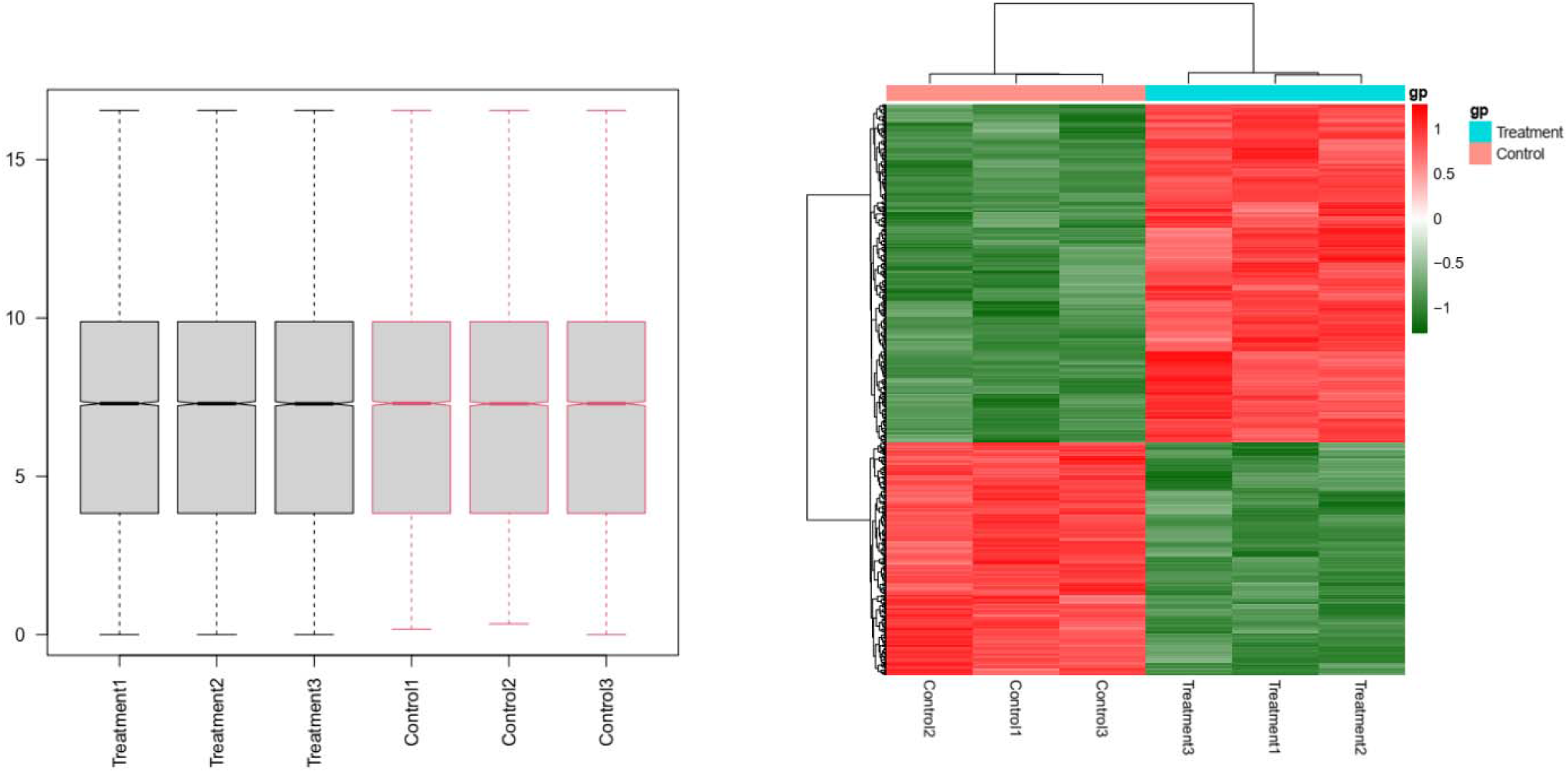
Heatmap in chloroquine-treated ESCC. Significant DEGs (P < 0.05) were used to construct the heatmap.

### Gene enrichments in chloroquine-treated ESCC

To study the features of significant genes, we introduced the gene enrichment analysis (Figure 2). We identified the top ten KEGG items, including “Cell cycle”, “Glycerophospholipid metabolism”, “Apelin signaling pathway”, “Glutamatergic synapse”, “p53 signaling pathway”, “Small cell lung cancer”, “Circadian entrainment”, “Pancreatic secretion”, “Pyruvate metabolism”, and “Base excision repair”. We also identified the top ten biological processes (BP), including “neuron projection extension”, “axon extension”, “response to unfolded protein”, “regulation of axon extension”, “positive regulation of axonogenesis”, “isoprenoid biosynthetic process”, “negative regulation of epidermal growth factor receptor signaling pathway”, “terpenoid biosynthetic process”, “pyrimidine deoxyribonucleotide catabolic process”, and “cellular response to follicle-stimulating hormone stimulus”.We then identified the top ten cellular components (CC), including “collagen-containing extracellular matrix”, “endoplasmic reticulum lumen”, “microtubule associated complex”, “vacuolar lumen”, “basement membrane”, “sarcoplasmic reticulum”, “inclusion body”, “sarcoplasm”, “chaperone complex”, “interstitial matrix”. We identified the top ten molecular functions (MF), including “nuclease activity”, “extracellular matrix structural constituent”, “endonuclease activity, active with either ribo-or deoxyribonucleic acids and producing 5’-phosphomonoesters”, “calcium-release channel activity”, “ligand-gated calcium channel activity”, “intracellular ligand-gated ion channel activity”, “proteoglycan binding”, “C4-dicarboxylate transmembrane transporter activity”, “amino acid:cation symporter activity”, and “DNA-(apurinic or apyrimidinic site) endonuclease activity”.

**Figure 2.**
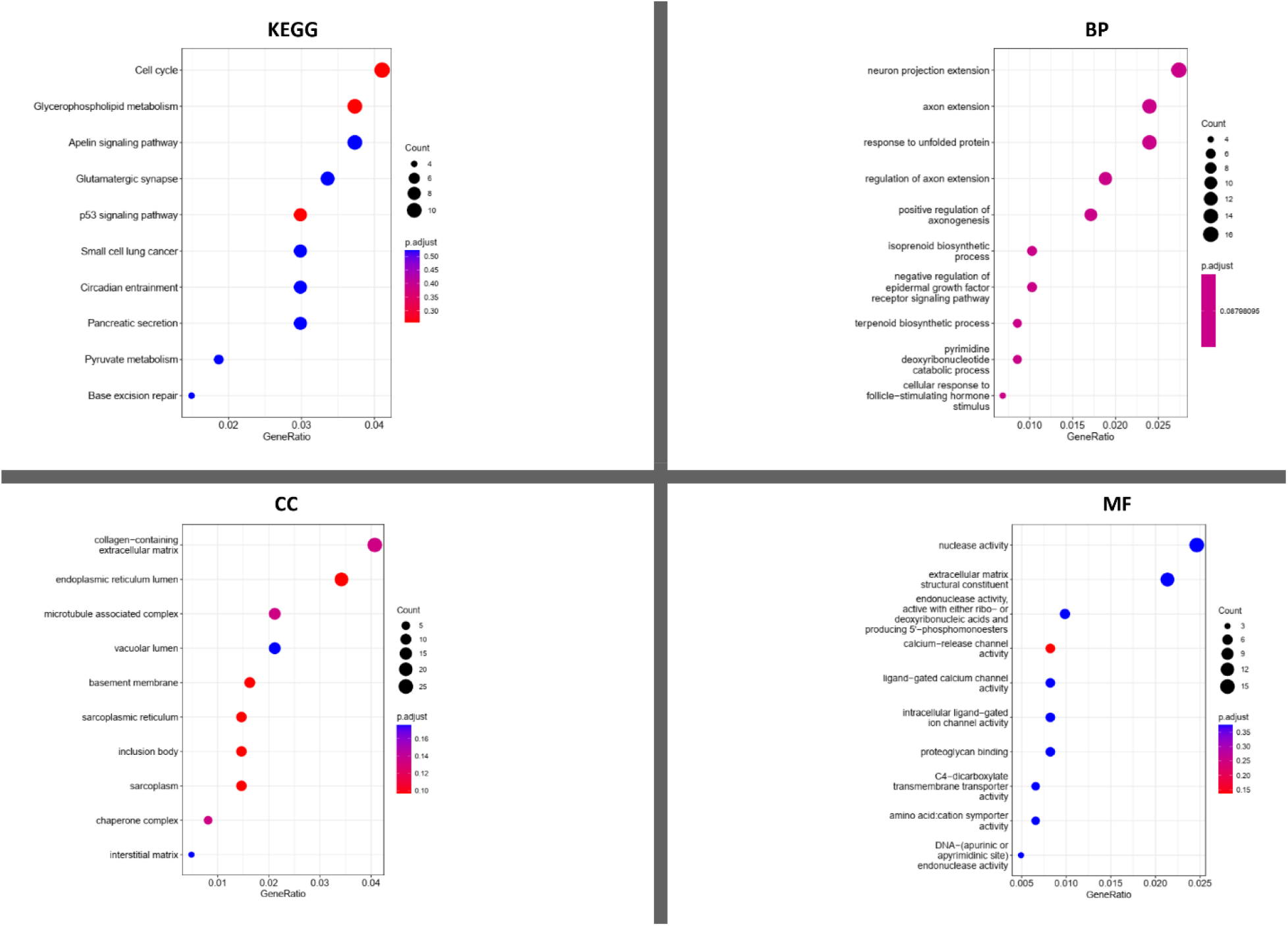
KEGG and GO analyses in chloroquine-treated ESCC. KEGG analysis; BP: Biological processes; CC: Cellular components; MF: Molecular functions.

### The protein-protein interaction (PPI) network and Reactome map analysis

The PPI network was created by using 643 nodes and 1461 edges. The top ten genes with the highest degree scores were presented in Table 2. We further constructed the top two clusters by analyzing the PPI network in Figure 3. Finally, we constructed the Reactome map by using the PPI network and DEGs (Figure 4) and we performed the Reactome analysis to further identify the top ten Reactome biological processes, including “HSF1-dependent transactivation”, “Attenuation phase”, “HSF1 activation”, “Regulation of HSF1-mediated heat shock response”, “Cellular response to heat stress”, “TP53 Regulates Transcription of Cell Death Genes”, “MET activates PTK2 signaling”, “Formation of Senescence-Associated Heterochromatin Foci (SAHF)”, “Transcriptional activation of p53 responsive genes”, and “Transcriptional activation of cell cycle inhibitor p21” (Supplemental Table S1).

**TABLE 2.**
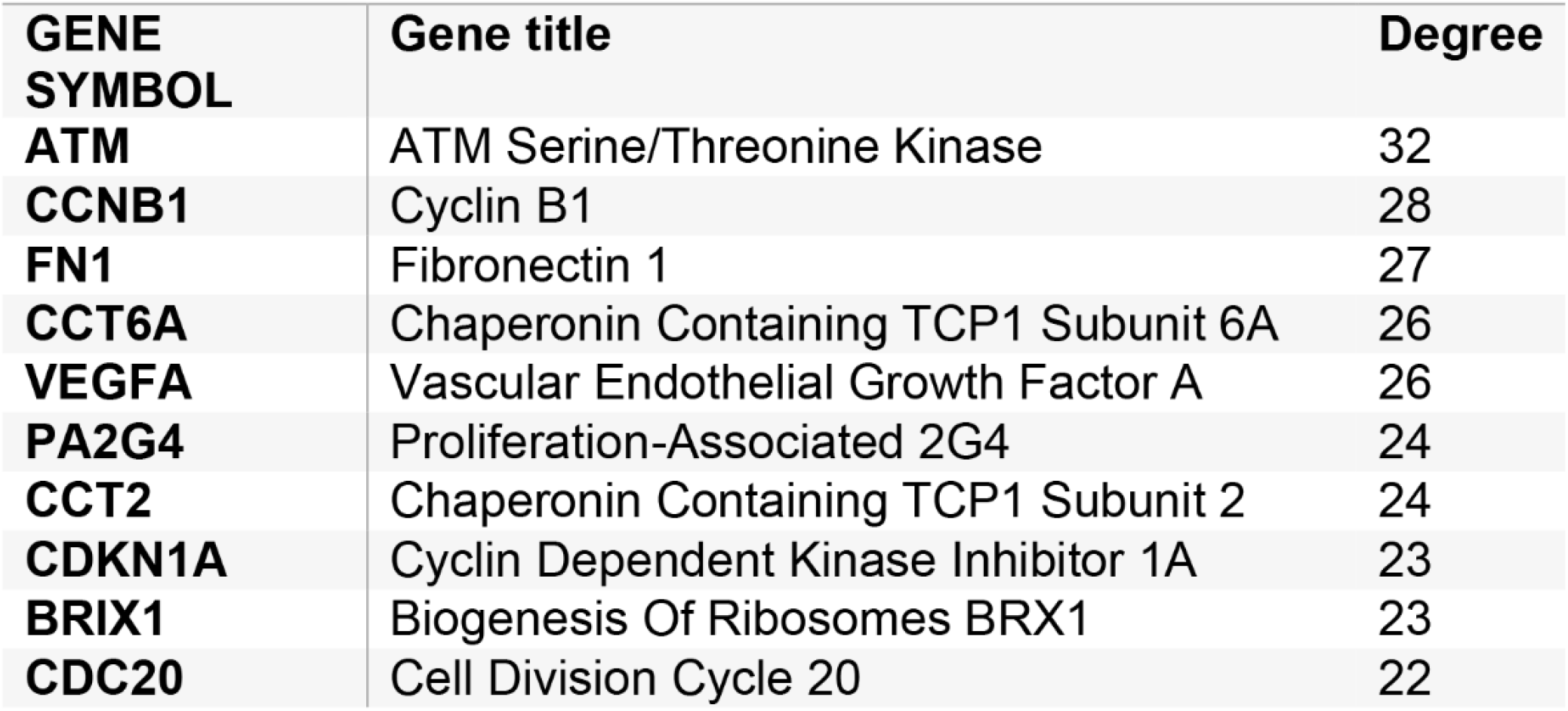
TOP TEN GENES DEMONSTRATED BY CONNECTIVITY DEGREE IN THE PPI NETWORK.

**Figure 3.**
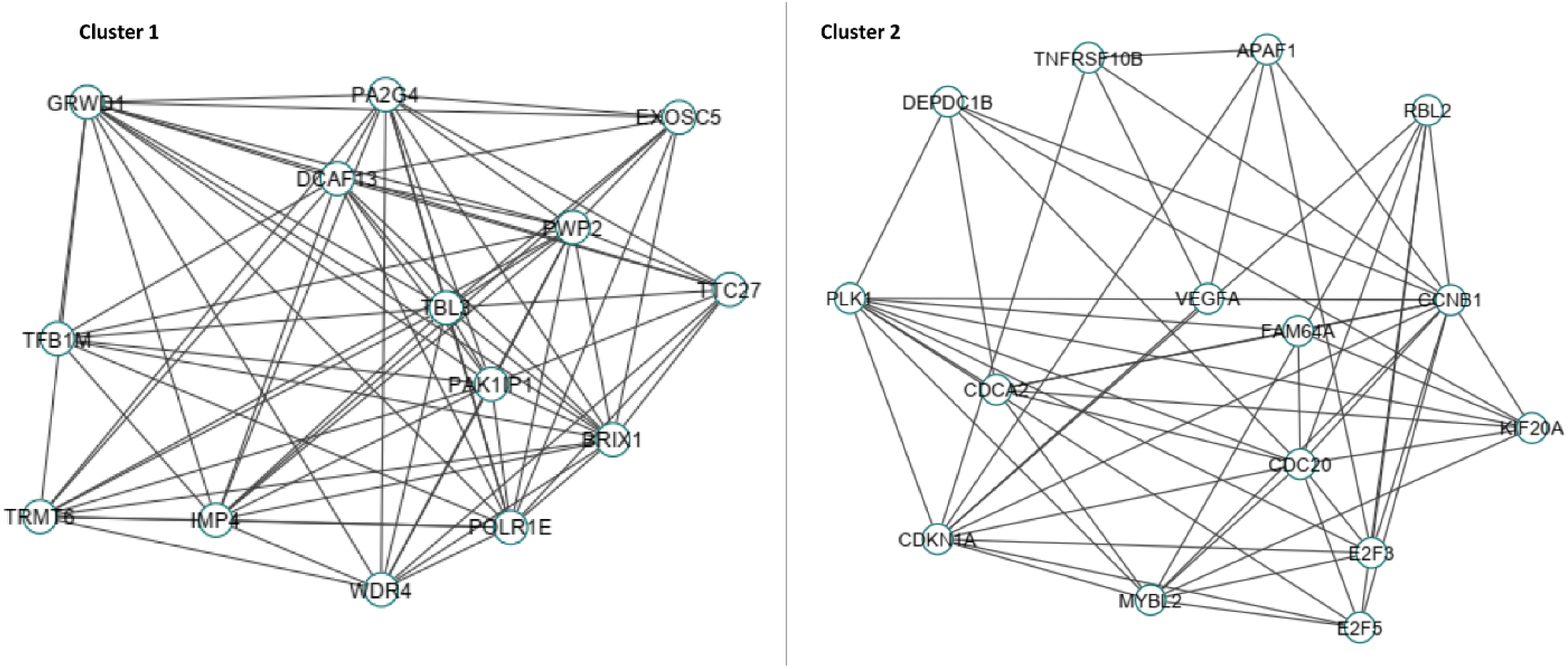
The PPI network analyses of DEGs in chloroquine-treated ESCC. The cluster 1 and cluster 2 were constructed by MCODE.

**Figure 4.**
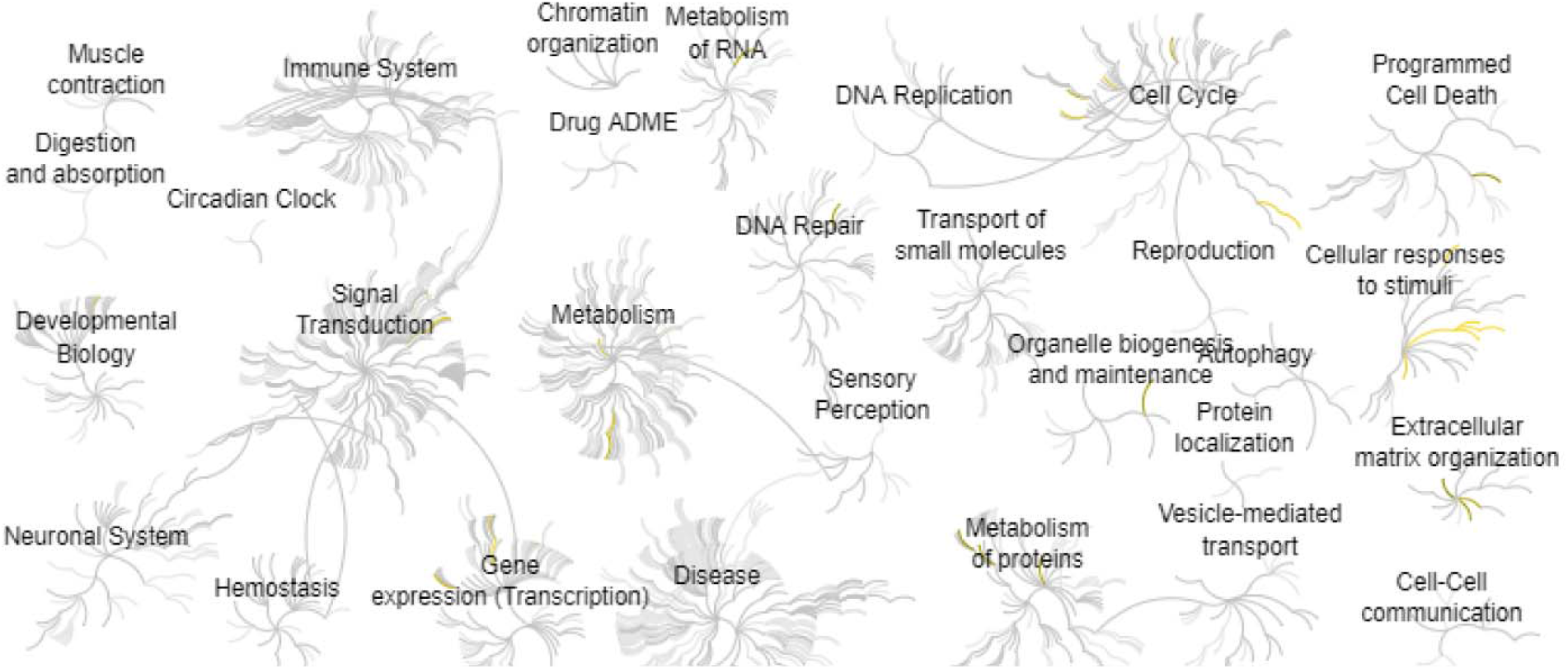
Reactome map representation of the significant biological processes in chloroquine-treated ESCC.

## Discussion

Chloroquine has the anti-tumor function by inhibition of autophagy, which causes the lysosomal membrane permeabilization (LMP) and contributes to apoptosis^15^. Esophageal squamous cell carcinoma (ESCC) is one of the human malignancies worldwide, but the mechanism of ESCC development is still unclear^16^. In this study, we analyzed the chloroquine-treated ESCC by using bioinformatic methods.

We identified the top two signaling pathways involved in the chloroquine-treated ESCC: Cell cycle and Glycerophospholipid metabolism. Interestingly, Keita Katsurahara et al found that ANO9 can mediate the cell cycle by increasing the number of cells in G0/G1 arrest in ESCC^17^. The study by Li-Yan Li et al found that SREBF1 controls the lipid metabolism and tumor-promoting pathways in squamous cancer^18^.

By constructing the PPI network, we identified the top ten interactive molecules that are associated with the chloroquine-treated ESCC. ATM and ATR kinases are the essential regulators in the DNA damage response (DDR), which serve as the apical mediators of the response to DNA double-strand breaks. Inhibition of DDR has been identified as an attractive treatment in cancer therapy. Thus, ATM kinase is a potential drug target for cancer treatment^19^. G Protein-Coupled Receptor signaling pathway involves a variety of pathological processes and diseases including arthritis, cancer, and aging^20-38^. Hanne Leysen et al also found the G Protein-Coupled Receptor Systems serve as critical regulators of DNA damage response processes by mediating ATM/ATR signaling^39^. Hui Zhang et al found the inhibition of CCNB1 showed decreased cell proliferation and S-phase cell proportion and increased apoptosis in pancreatic cancer^40^. The study by Lei Dou et al found the Chaperonin containing TCP1 subunit 3 (CCT3) expression was increased and the high levels of CCT3 further promoted the progression of cervical cancer by FN1^41^. The study by Guofen Zeng et al showed that the increased CCT6A can promote cancer cell proliferation by regulating the G1-to-S phase transition in hepatocellular carcinoma^42^. Lijun Liang et al found that the inhibition of autophagy repressed angiogenesis by VEGFA-related signaling pathway in non-small cell lung cancer cells^43^. Circadian genes are key transcriptional factors that involve a variety of cell functions including proliferation, differentiation, inflammation, migration, and secretion^44-55^. Elke Burgermeister et al found that BMAL1 is related to the bevacizumab resistance in colorectal cancers by regulating the VEGFA^56^. Yan Xu et al found the upregulation of PA2G4 is an unfavorable indicator in the progression of nasopharyngeal cancer^57^. TSPAN31 mediates the proliferation and apoptosis of gastric cancer cells by regulating the CCT2 signaling pathway^58^. CDKN1A has an important role in the response to the cisplatinlJpemetrexed combination in nonlJsmall cell lung cancer cells, which may be used as a predictive marker^59^. Jidanxin Ge et al foun the expression of BRX1 is closely related to malignant progression and prognosis of colorectal cancer^60^. The study by Lixia Wang et al found that CDC20 indicates an oncogenic function, which suggests that CDC20 can be a promising therapeutic target for cancer^61^.

In conclusion, our study identified the significant molecules and signaling during the chloroquine-treated ESCC. The Cell cycle and Glycerophospholipid metabolism are the major signaling pathways involved in the processes. Our study may provide a novel mechanism for ESCC treatment.

## Author Contributions

W.W, and Z. Q.: Conceptualization, Writing-Reviewing and Editing.

